# Regulation and function of trans-physeal growth plate bridges: evidence for a mechanical ‘base isolation’ role to minimise epiphyseal shear stress

**DOI:** 10.64898/2026.02.05.704004

**Authors:** D Valkani, LAE Evans, CM Disney, A Sharma, B Javaheri, J Chen, RT Hansen, M Hopkinson, S Monzem, P Louka, A Bodey, B Bay, PD Lee, KA Staines, AA Pitsillides

## Abstract

The cartilaginous growth plate (GP) is responsible for all bone elongation during post-natal growth yet must simultaneously contribute to mechanical epiphyseal stability for articulation. How the GP balances these dual functions across the bone-cartilage-bone epiphyseal interface during complex load-growth transitions is not defined. Herein, we examine regulation and mechanobiology of GP *bridges –* mineralised trans-physeal GP structures – to explore whether they serve these dual GP functions. We have determined the effects of age and sex, short- and long-term joint loading and several established and new osteotropic pharmacological agents on mouse tibial GP bridge number and areal density using micro-computed tomography. We also explored temporal formation and progression of GP bridges by serial in vivo scanning and we imaged epiphyseal load-transfer in young and mature mice in situ via synchrotron X-ray computed tomography (sCT) of intact joints under physiologically oriented load. Our utilisation of digital volume correlation revealed regional 3D load-induced strain inhomogeneities in the GP that are synchronised to bridge location and this was substantiated using finite element modelling. Furthermore, direct micro- and sCT examinations showed that bridges consistently contain a singular epiphyseal/metaphyseal discontinuity which appear to serve a novel mechanical ‘base isolation’ role to minimise shear stress across the GP. Our data indicate that bridges are regulatable, dynamic structures that synchronise GP strains and exhibit sensitivity to local joint mechanics. We highlight that trans-physeal bridges may contribute to longitudinal bone growth cessation whilst simultaneously stabilising the epiphysis by absorbing compression and shear strains.

## 1. Introduction

The growth plate (GP), a cartilaginous region sandwiched between the epiphysis and metaphysis in all bones containing a secondary ossification centre, is exclusively responsible for endochondral bone elongation until maturation (1, 2). The pace of this elongation shifts through life and its deceleration is primarily linked to slowed chondrocyte proliferation and changes in the velocity of senescence, suggesting that GP chondrocyte behaviour is dictated by overall growth rate (3-5). The GP must therefore serve as the source for all shifts in longitudinal bone expansion and yet must simultaneously contribute to preserving mechanical stability of the epiphysis during articulation. How the GP balances these dual and complex functions across this challenging bone-cartilage-bone interface is not defined.

Whilst these changing GP chondrocyte behaviours emphasise that growth history can influence GP function (5), the underlying mechanisms that enable the retention of GP structural integrity are not yet clear. Bone elongation is achieved by endochondral ossification processes in which a tightly regulated sequence of cartilage cell and linked matrix changes occur until the cessation of growth. Growth arrest heralds GP narrowing, as cartilage deposition slows and chondrocyte proliferation dwindles, culminating in a termination of longitudinal growth and epiphyseal fusion (6). This fusion ultimately involves replacement of the GP by a mineralised physeal ‘scar’ in most mammals, including humans (4, 7). However, how epiphyseal structural integrity is retained as the mechanical demands and varying loads from the adjacent articular joints are endured during bone elongation, remains relatively unexplored.

Epiphyseal fusion is often described to be rapid, as GPs in most mammals are typically defined as either entirely fused with no surviving chondrocyte columns, or *completely* unfused with well-preserved columns, implying speedy replacement of all chondrocytes, *presumably* by vascular and bone cells. Timing of fusion varies by anatomical position, suggesting additional local control (6, 7). Fusion is an active process governed by its own hormonal controls, cell mechanisms and structural features that – whilst certainly preceded by – do not automatically follow growth cessation (6). Thus, timing of fusion can be advanced or delayed in specific disorders (7, 8); e.g. estrogen deficiency preserves an unfused epiphysis long after growth has stopped, whereas its substitution results in a resumption of the normal fusion timetable (5-7). These points have clinical impact as nonsurgical options to manipulate adult height and anisomelia are limited by epiphyseal fusion. Vitally, these clinical options are further limited as the mechanisms controlling epiphyseal fusion remain to be completely resolved.

One reason for this lack of understanding is that our knowledge of GP fusion is mostly based on animal studies where interspecific variation is inherent; while humans and rabbits acquire fully fused GPs that are completely infiltrated by trans-physeal mineral, rats and mice do not. This has obscured how data derived from rodents should be interpreted (8). Micro-computed tomography (µCT) images of rat tibiae indeed show that maturation in rodents is instead marked by an accumulation of mineralized struts, termed *tethers* or *bone bridges* (bridges hereafter) which traverse this trans-physeal bone-cartilage-bone GP interface (9). These bridges have been documented in mouse, rat, dog, pig and human GPs (9-13), where they are reported to perpendicularly connect epiphyseal and metaphyseal bone surfaces. Although rat and murine GPs never completely mineralise to fully fuse, the bridges are reported to accumulate, especially in adolescence as GP thickness decreases (9, 10). The function of these bridges and their regulation is, however, almost entirely undefined and the factors controlling their number and distribution, and their relationship to bone growth, is poorly appreciated.

Our µCT or synchrotron X-ray CT (sCT) imaging studies have established a protocol for quantifying bony bridges, their distribution and local areal density in 3D across the entire murine epiphysis (14-16). This was used to reveal that osteoarthritis-prone STR/Ort and healthy CBA mice both display overt bridges prior to growth cessation, and that spatial bridge clustering diverges in these mouse strains, suggesting that their emergence and patterning is a controlled process (14). This notion that bridge formation is regulated is supported by elegant studies showing that vitamin D receptor knockout mice have fewer bridges with altered distribution, and by data linking these shifts to GP morphology and weight (10, 17). Whether bridges are regulated by pharmacological osteotropic agents or GP mechanics is, however, poorly appreciated, as is their role, if any, in local mechanics or long bone growth.

While bridge function remains somewhat mysterious, it has been speculated that they stabilise the GP thereby serving a protective mechanical role during maturation (9, 17). This has yet to be examined directly, however, and it is now possible using images captured by sCT during in situ joint loading to quantify such strain fields volumetrically by digital correlation methods. This enables strains to be measured with nanoscale precision in the GP and mapped specifically to bridge location (16).

Herein, we evaluate the effects of age and sex, the impact of joint loading, and several established and new osteotropic pharmacological agents on mouse GP bridges. We explore temporal bridge evolution using serial in vivo scanning, and image load-transfer through epiphyses using sCT in intact joints during physiologically oriented in situ loading to expose ground-breaking shifts in GP deformation in growing and skeletally mature mice. Nanoscale 3D mapping of these strain environments reveals regional inhomogeneities that are synchronised to mineralised bridge location, which we substantiate by finite element simulation. Finally, our imaging of intact GPs reveals the consistent presence of an intra-physeal mineral *discontinuity* in each bridge. These data indicate that bridges are regulatable, dynamic structures that synchronise GP strain environments whilst also exhibiting sensitivity to local joint mechanics. They highlight a new paradigm, where intra-physeal mineral discontinuity within bridges can simultaneously prolong longitudinal GP bone expansion function whilst also serving as a novel shock absorber to minimise load-induced stress during GP maturation.

## 2. Materials and Methods

### 2.1 Mouse colonies

C57Bl/6J, CBA and STR/Ort mouse colonies were established in the Biological Service Unit, Royal Veterinary College with appropriate breeding programs, housed in conventional polypropylene cages maintained in regulated humidity, between 19–23°C with 12-hour light/dark cycles. Mice were fed ad libitum with standard RM1 maintenance diet (Special Diet Services, South Witham, UK) and water. Schedule 1 methods were used for euthanasia. All procedures were accord with the Animals (Scientific Procedures) Act 1986, and were approved by the Royal Veterinary College Ethical Review Committee and the United Kingdom Government Home Office under specific licence and ARRIVE guidelines.

### 2.2 Experimental design

#### 2.2.1 Overall approach

Bridge number was evaluated in proximal tibial GP scans from mice (Supp. Table 1), including in: i) different sexes, ii) right and left limbs to assess symmetry, iii) lateral and medial compartments to asses relationship with predominant habitual loading, iv) distinct ages (young/mature/old), v) diverse strains (C57Bl/6J, CBA, STR/Ort), vi) the presence or absence of pharmacological osteotropic factors (PTH, SFX-01) and, vii) in mice in which the right knee joint was subjected to controlled mechanical loading (left, contralateral unloaded control).

#### 2.2.2 Mechanical loading

Right knee joints were loaded in vivo under isoflurane anaesthesia in male and female mice (strain as specified) at specific ages. Mechanical loading was achieved using the axial compression model of Souza et al (18), and depending on study objective different load regimens were used; these differed either in load period duration (single/double episodes, or 6 episodes over a 2-week-long period) and applied load magnitude (9-12 N) (19). In most cases, each load episode involved application of a defined dynamic axial load of 9-12 N, with 2 N hold load (0.1 sec trapezoidal-shaped pulse period, with 0.025 sec fall time, 0.05 sec hold, 0.025 sec rise time, 9.9 sec rest), 10 sec each cycle, 40 cycles) applied to the right limb. At study termination, left (contralateral) and right (loaded) hind limbs were dissected, excess muscle and tissue removed, fixed in 4% paraformaldehyde (PFA) for 24 hours before a brief wash and storage in 70% ethanol.

#### 2.2.3 Mechanical bone loading with parathyroid hormone (PTH) treatment

Weight-matched 12-week-old C57Bl/6J female mice received either an interrupted (5 days on/2 days off) regime of 40 µg/kg PTH (1-34; 8 µg/ml) morning injections or matching injections of vehicle for three weeks (n=9/10 per group; Charles River/Harlan); administration was performed after the anaesthesia on days when the tibia was loaded (20). After one (first) week of dosing, the right hind limb of each mouse was subjected to 12 N dynamic axial loading under anaesthesia, for 3 days each week, for 2 weeks (as above); the left hindlimb served as a non-loaded contralateral control (20).

#### 2.2.4 Sulforadex^tm^ (SFX-01) treatment

Two strains (C57Bl/6J and STR/Ort) were used. Groups receiving SFX-01 were treated with ∼100 mg/kg SFX-01 in drinking water (vehicle, water) which was refreshed every other day SFX-01 (21, 22). Mouse weight was recorded from baseline and every week for the duration of the study. In the first study, 9-week-old male C57Bl/6J mice were assigned randomly into groups (n=6-8/group) and treated with SFX-01 in drinking water (or water ad lib) for 3 weeks and all mice were euthanised on the first day of week 4. In the second study, 14-week-old male STR/Ort mice (n=11/group) were treated similarly with SFX-01 in drinking water for 5 months (with controls receiving water ad libitum).

#### 2.2.5 Tamoxifen administration

Seven week-old male C57Bl/6J mice (Jackson Laboratory, USA) were treated with tamoxifen or vehicle alone as described by Lo Cascio et al. (23) Briefly, 20 mg/ml tamoxifen was dissolved by sonication in 90% sunflower oil and 10% EtOH and was administered at 50-100 mg/kg via IP injections on three consecutive days. Mice were sacrificed two weeks later.

### 2.3 Scanning and visualization

#### 2.3.1 Benchtop µCT scanning for serial in vivo imaging of bridges

The right tibia of 12-week-old female C57Bl/6J mice was subjected to loading (6 episodes of load, each with 40 cycles, 12 N at 2 Hz, 10 s cycle time, as above) under isoflurane-induced anaesthesia over a 2-week-long period and multiple in vivo µCT scans were collected on experimental days 0 and 14 (coinciding with timepoints before and immediately after 2-week-long period of loading) and day 101 (corresponding with 12 weeks post-loading) (23). In vivo entire right tibia scans were achieved using the SkyScan 1176 (Kontich, Belgium), with X-ray tube operated at 40 kV and 600 μA, orbit of 198°, 1 mm aluminium filter, 0.8° rotation angle and 9 μm voxel size (∼0.5 hour per sample) (24). Right and left tibiae of all mice were ex vivo scanned using a Skyscan 1172 (Bruker, USA); X-ray tube at 50 kV and 200 μA, 960 ms exposure time with a 0.5 mm aluminium filter, 0.6° rotation angle at voxel size of 5 μm.

#### 2.3.2 CT scanning for bridging analysis

sCT was performed on the I13-2 beamline (Diamond Light Source, Oxfordshire, UK; Beamtimes MG24786-2 and MG28353-1) and in all cases the experimental unit was the knee joint from an individual mouse which had been dissected, skinned, with most surrounding musculature removed. The proximal tibia and distal femur had been cut through the metaphysis following extraction such that the articular surface and GP regions remained intact. The knee joints were wrapped in 1x PBS-soaked gauze, fresh frozen and stored at −20 °C until use.

Samples were scanned at room temperature or at −20°C in a bespoke cold stage (16). Images were acquired using a filtered pink beam with an average energy of 19 keV (based on an energy sensitivity study for optimum image contrast). 4,001 projections per sample were using an exposure time of 0.12 seconds during 360° rotation and were detected used a PCO 4000 CCD imaging camera with 4x objective lens (total magnification 8x) at an effective voxel size 1.125 μm^3^. The field of view (FOV) was 4.5 mm x 3 mm and scan time was 1.1 minutes. Projections underwent darkfield and flatfield correction, ring artefact removal and distortion correction prior to reconstruction by filtered back-projection in the Data Analysis Workbench (DAWN) software into 8-bit.tiff stacks (25-27).

#### 2.3.3 sCT scanning for digital volume correlation analysis (DVC)

sCT was performed on whole mouse knee joints under realistic load conditions using the I13-2 beamline as previously described (16). Briefly, intact knee joints were mounted between two specialized, 3D printed plastic cups within the P2R in situ mechanical loading rig in preparation for axial compression (28). sCT projections were acquired using a filtered pink beam (average energy of 5-30 keV; 1.3 mm carbon, 3.2 mm aluminium, 0.75 mm silver) with an undulator gap of 5 mm. A total of 4001 projections were collected for each scan over 180° of continuous rotation (fly scan), with a 20-40% transmission, 4.1 x 3.45 mm field-of-view and an effective pixel size of 1.6 or 0.8 µm depending on the setup. Images were collected by a PCO.edge 5.5 (PCO AG, Germany) detector (16-bit, 100 fps) mounted on a visible light microscope of variable magnification. Radiation dose was approximated to be between 100 kGy to 240 kGy.

##### 2.3.3.1 DVC analysis

To resolve in situ growth plate strains by DVC, we first upgraded a regime for recapitulating knee joint loading in intact murine hindlimbs during typical locomotion compatible with sCT imaging (19). In line with previous studies, an initial 2 N load was applied to fix hindlimbs in a flexed positionand as anticipated, joints exhibited viscoelastic stress relaxation under this 2 N ‘baseline’ uniaxial load and to prevent motion artifacts, 15 minutes was allowed after load application (<4 N) prior to sCT imaging, when force stabilisation was attained (19). Successive sCT images were captured at displacement-controlled load steps All samples were wrapped in parafilm to stop them drying out performed and placed in the P2R rig.

### 2.4 Bone analysis and segmentation for bridging analysis

Raw µCT images were reconstructed on NRecon®, reoriented in DataViewer® and segmented using CTAn® (Bruker, Belgium), with femur, fabellae, fibula, patella and all bones of the foot removed by drawing around the tibia and a new segmented region of interest saved (16).

#### 2.4.1 Growth plate bridging analysis

GP bridge analysis was performed on tibial scans, obtained either from benchtop µCT or sCT imaging, using Avizo® (Thermo Fisher Scientific, USA) software (15, 29). Briefly, the central points of all bridges (Fig. 1A) were identified and projected onto the tibial joint surface (Fig. 1B) and bridge number and areal density were calculated in each knee joint. 3D reconstructions of each joint, including the GP (Fig. 1C), were generated in Avizo, and each bridge, which crosses the GP is allocated a heat-map colour that represents the areal number density at each bridge location (Fig. 1B, left). Visualisation and colour-coding of bridge number and areal density was performed in Avizo, Image J and CTVox® 3.1.0 r1167 version (Skyscan, Kontich, Belgium). For some joint region-based analysis, the GP was divided into lateral and medial compartments (Fig. 1B, middle and right).

**Figure 1.**
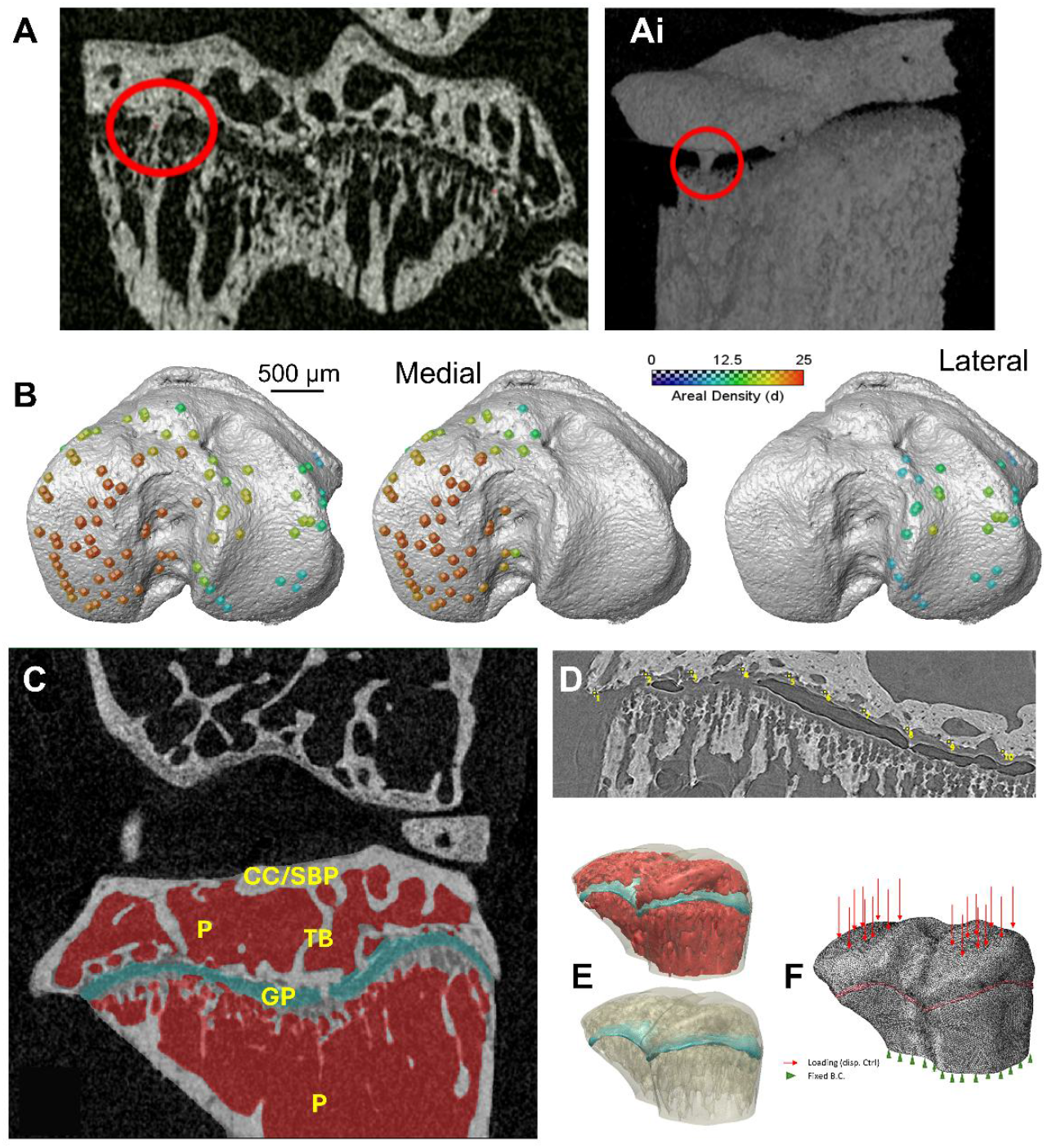
Visualisation of growth plate (GP) bridges by Avizo (A, left) and CTVox (Ai, right). Bridge location/areal density across 14-week-old C57Bl/6J GP projected onto tibial articulation (B, left), split to show medial (middle) and lateral condyle (right) bridges. Representative µCT slice (C) segmented to identify GP (blue), calcified cartilage (CC), trabecular bone (TB), subchondral bone plate (SBP) and porous marrow space (P, red). Point cloud collection on epiphyseal side (D). 3D render of segmented models to show epiphyseal porous space (upper) and TB (lower) and locations relative to GP (E). Image-based FE modelling of biomechanical bridge properties in GP with uniform load conditions assigned to two loading areas of CC/SBP in both medial and lateral compartments (F).

### 2.5 Data processing and DVC analysis

#### 2.5.1 Quantification of strain at growth plate bridges

Projections were reconstructed with dark- and flat-field correction and ring artefact suppression prior to filtered back-projection using the Diamond Light Source software DAWN (26, 30). The joint scans were cropped, normalised and 3D median filtered (using a 2-kernel size). In order to identify the GP deformation induced by loading in young, skeletally mature and ageing mice, sCT images stacks were resliced in Fiji/Image J and a set of points in a series of scans (20 points from each of 20-25 scans for each raw file) were manually placed across the epiphysis and metaphysis, with point coordinates exported in.csv format and utilised for point cloud generation (Fig. 1D). Each point specifies the location at which the point cloud will be placed to measure displacement by DVC. In each scan, ‘point clouds’ (initial x, y, z positions) in two groups, one on the epiphyseal and one on the metaphyseal side of the GP, were defined using MATLAB; manually placed points from epiphyseal and metaphyseal surfaces were smoothed using locally weighted regression to create continuous surface fits, and a regular grid of 8 pixel spacing was sampled and trimmed to the original data footprint to create the DVC point cloud.

As described by Madi (16), two 8-bit image volumes (non-deformed/deformed) were used as input to the DVC open-source code supported by CCPi (Collaborative Computational Project in Tomographic Imaging, https://tomographicimaging.github.io/iDVC/). DVC code was then used to calculate the displacements at each point, which defined the location of spherical subvolumes (30 voxels, 5000 sampling points) for correlation using zero-normalized sum of squared differences (ZNSSD) as the objective function and tricubic interpolation (30-32). A maximum displacement limit (10 pixels, 12 degrees of freedom) was applied during correlation, with initial rigid transformation applied. Matlab R2019a (Mathworks Inc., Natick, Massachusetts, USA) was used for post DVC processing in preparation for finite element-based strain calculation (31). Surface normals were calculated for the metaphyseal surface and used to guide epiphyseal node placement, producing a well-aligned mesh between the two regions. Hexahedral elements were formed by pairing adjacent metaphyseal nodes with corresponding epiphyseal nodes, and strain was computed at element centres using nodal displacement data, followed by extraction of principal strains and directions.

For measurement and examination of strain around the location of GP bridges, Image J was used to locate the bridges on the reconstructed and resliced image volumes. Bridges were manually selected and saved in a file (.csv), producing a file with coordinates x, y, z, similar to that produced for point cloud identification; these were used to project specified bridges onto DVC strain maps. Specifically, to quantify strain at each anatomical bridge location, a nearest-neighbour approach was applied using the 3D strain calculation points as a reference. For each predefined bridge point, the ten closest strain points were identified, and their strain values were averaged to represent local strain behaviour.

#### 2.5.2 Quantification of strain orientation across the growth plate

Scans from non-loaded joints served as a reference for initial GP point cloud selection and for tracking, using datasets from sequential scans captured with progressive load application (<4 N). Spatial registration of datasets was performed in MATLAB ahead of point cloud generation. Strain orientation was calculated for epiphyseal and metaphyseal compartments and combined to extract respective strain at all GP locations (Fig. 1D); allows DVC point cloud vector mapping (between epiphyseal and metaphyseal surfaces intersecting normal metaphyseal vectors) to lateral-medial/posterior-anterior GP architecture and 3D GP bridge position. To delineate engendered forces, total strain (top-bottom), maximum shear, tension and compression were assessed. Deformation point clouds on combined epiphyseal/metaphyseal parts were used to extract overall strains, and these were used to generate histograms to expose cumulative incidence across the entire GP.

### 2.6 Image-based finite element modelling

Image-based finite element (FE) models were developed to assess the biomechanical effects of bridges in the GP. µCT images were segmented using Synopsys Simpleware (Version U-2022.12-SP2, Synopsys Inc.) to distinguish three phases based on distinct X-ray attenuation characteristics: the GP, mineralized structures (calcified cartilage (CC), subchondral bone plate (SBP), trabecular bone (TB) and GP bridges (B)), and the porous trabecular space (P; Fig. 1C). FE models were generated from segmented images and meshed using C3D4 elements with adaptive meshing control for improved accuracy. Uniform compressive loads of 1 MPa were applied to the main loading areas on lateral and medial condyles, simulating load transmission through the hyaline cartilage, while the model’s distal end was fully constrained (Fig. 1E-F). To isolate geometric bridge effects, the linear elastic, isotropic and homogenous properties were assumed (33-36). Von Mises stress as one of key biomechanical estimates (37, 38) was evaluated within the mineralized structures and the GP.

### 2.7 Statistical analysis

For statistical analysis, the GraphPad Prism® 9 software was used. Data are presented as mean ± SD. For two variable comparisons, t-test or Mann-Whitney test were used. For more than two variables one-way ANOVA was used. P values less than 0.05 (p ≤ 0.05) were considered statistically significant.

## 3. Results

### 3.1 GP bridge number is greater in males and escalates with age in both sexes

We first quantified the total number of bridges in the tibial GP in matched left-right limbs, and in mice of both sexes across a range of ages in divergent strains (C57Bl/6J, unless otherwise stated). These analyses showed that total bridge number in right and left limbs of the same mice showed tight symmetry with similar number of GP bridges in 14-week-old male mice (Fig. 2A). This establishes that contralateral limbs are likely suitable as controls in situations with ipsilateral intervention, and that GP bridge number is likely a controlled process. We also found that tibial GP bridge number was significantly greater in males than in age-matched female mice at 14-weeks-of-age (p<0.05), when longitudinal growth is deemed close to cessation (Fig. 2B). Quantification also revealed that this sexual dimorphism in bridge number became more marked as mice reached 18-22-weeks-of age (p<0.001), with female GPs containing <50% fewer bridges (vs. males; Fig. 2C).

**Figure 2.**
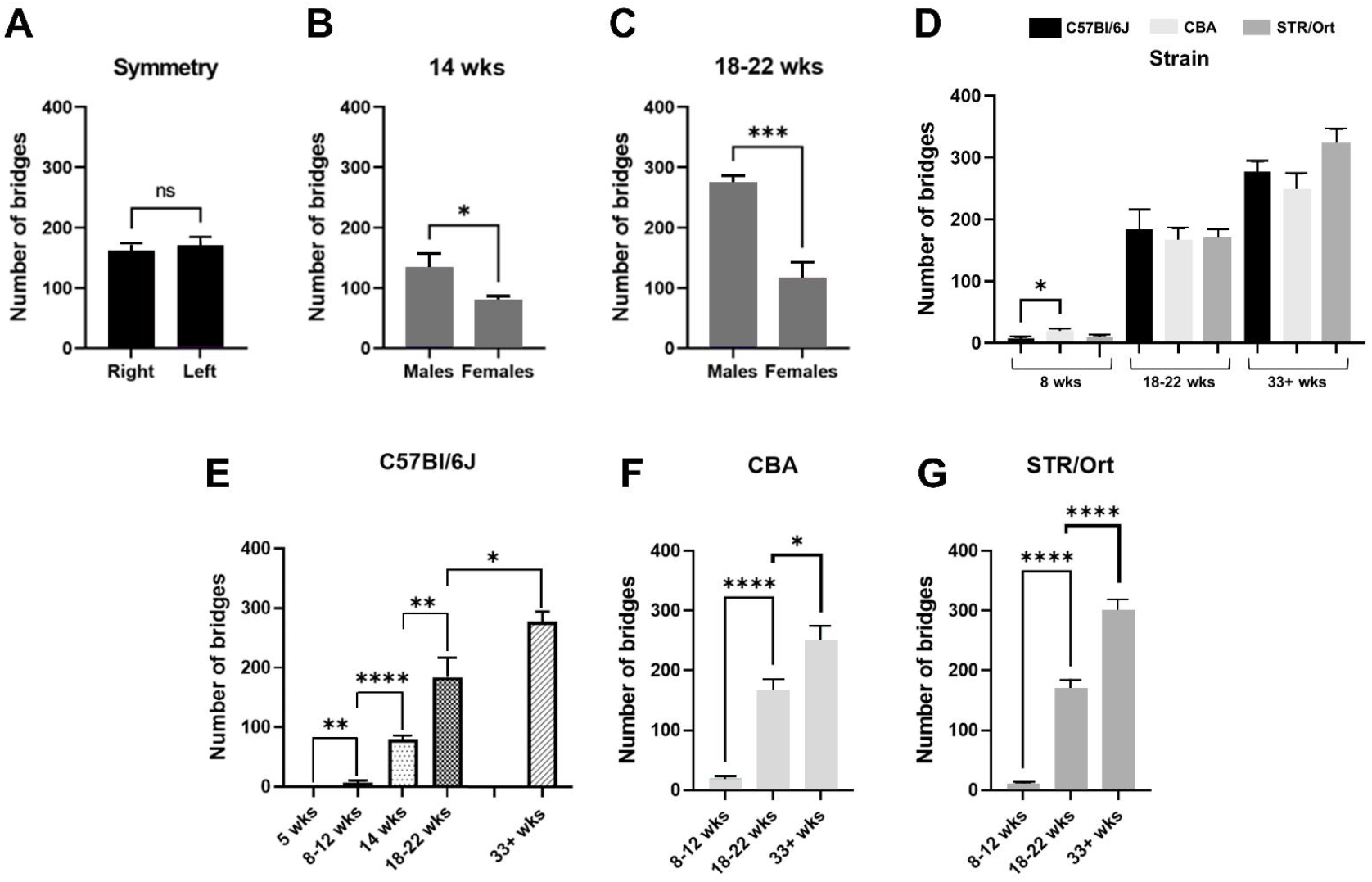
Bridge number exhibits left-right symmetry in tibiae of 14 week-old male (A) and sexual dimorphism (fewer in females) in both 14 week-old (B) and 18-22 week-old (C) C57Bl/6J mice. There are only minor differences in the total number of bridges in age-matched male mice of three different mouse strains (C57Bl/6J, CBA and STR/Ort) with significantly more bridges in CBA than C57Bl/6J at 8 weeks of age (D). Total bridge number increases and accrual rate decreases with age after 22 weeks in male C57Bl/6J (E), CBA (F) and STR/Ort mice (G). *p<0.05, **p<0.005, ***p<0.0005, and ****p<0.0001.

To interrogate age-related accumulation further, GP bridge number was quantified across a wider age range in male C57Bl/6J mice. This revealed that greatest bridge accumulation occurred in male mice between 12 and 14-weeks-of-age (p<0.0001), with further age-related increases in bridge number occurring less rapidly (Fig. 2E). As bone mass likewise increases in mice across this age range, we explored possible links between genetically-encoded bone mass differences and bridge numbers in C57Bl/6J, CBA and STR/Ort strains; chosen as they have been shown to possess divergent basal bone mass phenotypes (C57Bl/6J< CBA< STR/Ort) (39), are from both close (CBA and STR/Ort) and distant genetic lineages (C57Bl/6J) and, incidentally, show divergent osteoarthritis predisposition (STR/Ort>C57Bl/6J>CBA) (40). This exposed a relatively conserved age-related accumulation of bridges in all three mouse strains, with fastest rates from 8-12 to 18-22 weeks and significantly greater numbers at 33+ weeks of age (vs. 18-22 weeks; Fig. 2D-G). Small shifts in this age-related bridge accumulation pattern were seen in STR/Ort in which there were significantly greater numbers (p<0.0001) at 33+ week of age than in C57Bl/6J (p<0.005) and CBA (p<0.05) mice, and in CBA at the youngest age (p<0.05; Fig. 2D), indicating that skeletal maturation and bridge formation are linked and accumulation rates are broadly similar in strains with markedly divergent bone mass.

### 3.2 GP bridging is regulated by osteotropic pharmacological agents and by mechanical loading

To assess whether GP bridging exhibits sensitivity to pharmacologic agents capable of regulating bone mass, we also measured bridge number after treatment with osteotropic agents, including: i) a regime of intermittent PTH (1-34) known to produce increases in cortical and trabecular bone mass (20), ii) tamoxifen, a synthetic ‘anti-estrogen’ with effects on bone mass and density (40, 41) or iii) SFX-01, a novel stable form of sulforaphane that increases bone mass largely via anti-osteoclastogenic actions (20, 42, 43) on the basis that this would inform likely mechanism of bridge regulation. We found bridge number to be unmodified (vs. vehicle) in the tibial GP of 11-week-old female mice treated with PTH (Fig. 3A). In contrast, week-long tamoxifen treatment of 11-week-old male mice yielded small yet significant (p<0.05) decreases in bridge number (vs. vehicle; Fig. 3B), suggesting links between accumulation/reduction of bridges and regulation of longitudinal bone growth. Interestingly, the GP in 14-week-old male STR/Ort mice treated with the novel anti-resorptive SFX-01 showed significantly greater bridge numbers (vs. vehicle; p<0.0001; Fig. 3C); this accumulation of bridges with SFX-01 was however, absent in older mice (Fig. 3D), suggesting that SFX-01 effects on GP bridging are restricted to young mice, linking a pharmacologic anti-resorptive and the control of bridge accumulation.

**Figure 3.**
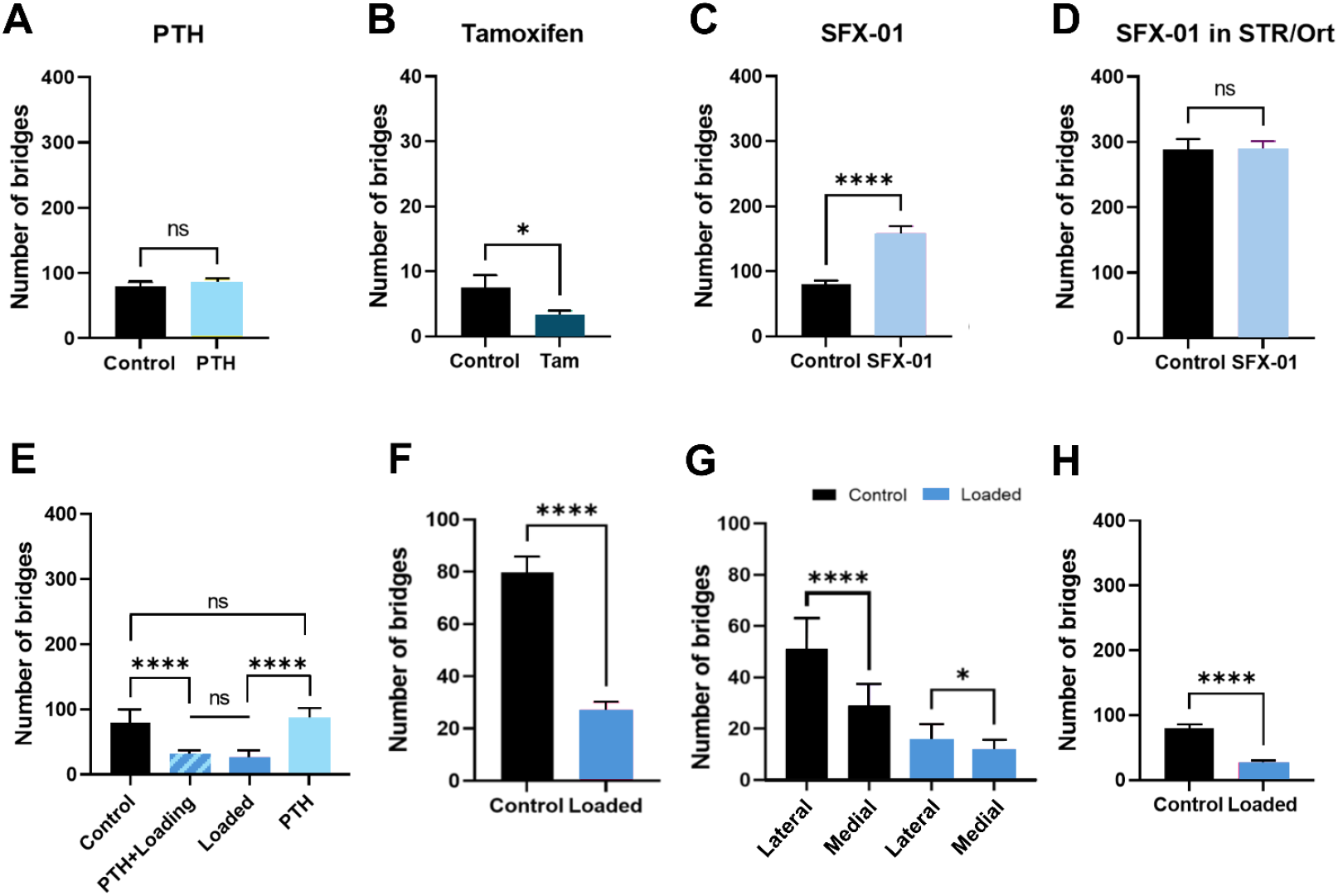
PTH treatment for 3 weeks does not alter bridge number in female C57Bl/6J mice aged 12 weeks (A), tamoxifen treatment for 3 days decreases bridge number in male C57Bl/6J aged 7 weeks (B) and SFX-01 treatment for 3 weeks increases bridge number in male C57Bl/6J mice aged 9 weeks (C) but does not change bridge number after 5 months in 14-week-old STR/Ort mice (D). Evaluation of number of bridges in 15-week-old female C57Bl/6J mice after 3 weeks of PTH administration alone or in combination with 12N load (E) shows that PTH does not alter but loading reduces bridge number. Loading at lower 9N magnitude also significantly decreases bridge number in 14-week-old female (F) and male C57BL/6J mice (H), with greater reduction in lateral than medial compartment in females (G). *p<0.05, **p<0.005, ***p<0.0005, ****p<0.0001.

It is known that PTH can act either additively or synergistically with load to regulate bone mass (20, 44). To examine whether similar interaction is seen in bridge regulation, their number was measured in mice treated for three weeks with intermittent PTH during which the right tibia (non-loaded control; left tibia) was exposed to a 2-week long 12 N mechanical loading protocol with established bone accrual effects (20). This verified that exogenous PTH did not modify the number of bridges and additionally that load and PTH did not interact to regulate bridge number (Fig. 3E). Intriguingly, it was found that mechanical loading of the tibial epiphysis generated a significant reduction in GP bridge numbers disclosing a novel regulation of bridge formation by the local mechanical milieu (Fig. 3F).

To interrogate this mechanical restraint on bridge formation, we exploited our prior work which found that lower loads exerted only minimal, if any, osteotrophic effects and that greatest deformation in this loading model are engendered latero-posteriorly (18, 19). We thus performed similar studies at lower 9 N magnitudes in which bridge number was evaluated across defined anatomical regions to identify potential regionalisation in their numerical reduction. This showed significantly more bridges in the lateral (n>40) than medial (n<40) compartment in control non-loaded limbs of female mice and, vitally, that 2 weeks of 9 N loading significantly reduced bridge number (p<0.0001, vs. non-loaded, Fig. 3G). Compellingly, this constraint on bridge number was intimately linked to applied local loading as there was a greater reduction in bridge number in the lateral (67% fewer; p<0.05) than medial compartment (50% fewer; p<0.05; Fig. 3G); bridge number was also significantly reduced by 9N loading in male C57BL/6J (Fig. 3H), supporting a sexually-conserved effect of joint loading.

### 3.3 GP bridges appear unrelated to growth direction and sequential 3D µCT evaluation confirms their regionally-matched, temporary deficiency in response to applied load

High resolution imaging was performed to inform views on GP bridge origins, formation mechanism and any potential directionality in their traversing of the GP. Visualisation of bridges by sCT (or µCT) failed to disclose resident osteocytes and instead suggested that they may be composed of calcified cartilage (Fig. 4A-D). Inspection of bridges revealed that there was unlikely a singular origin, with incomplete bridges equally likely to be connected to epiphyseal or metaphyseal GP face (Fig. 4A-D). We also measured bridge widths to find that they exhibited a >5-fold range in diameter, with cursory sub-grouping suggesting that the widest bridges are rarest, implying that their trajectory of emergence is faster than their subsequent thickening (Supp. Table 2).

**Figure 4.**
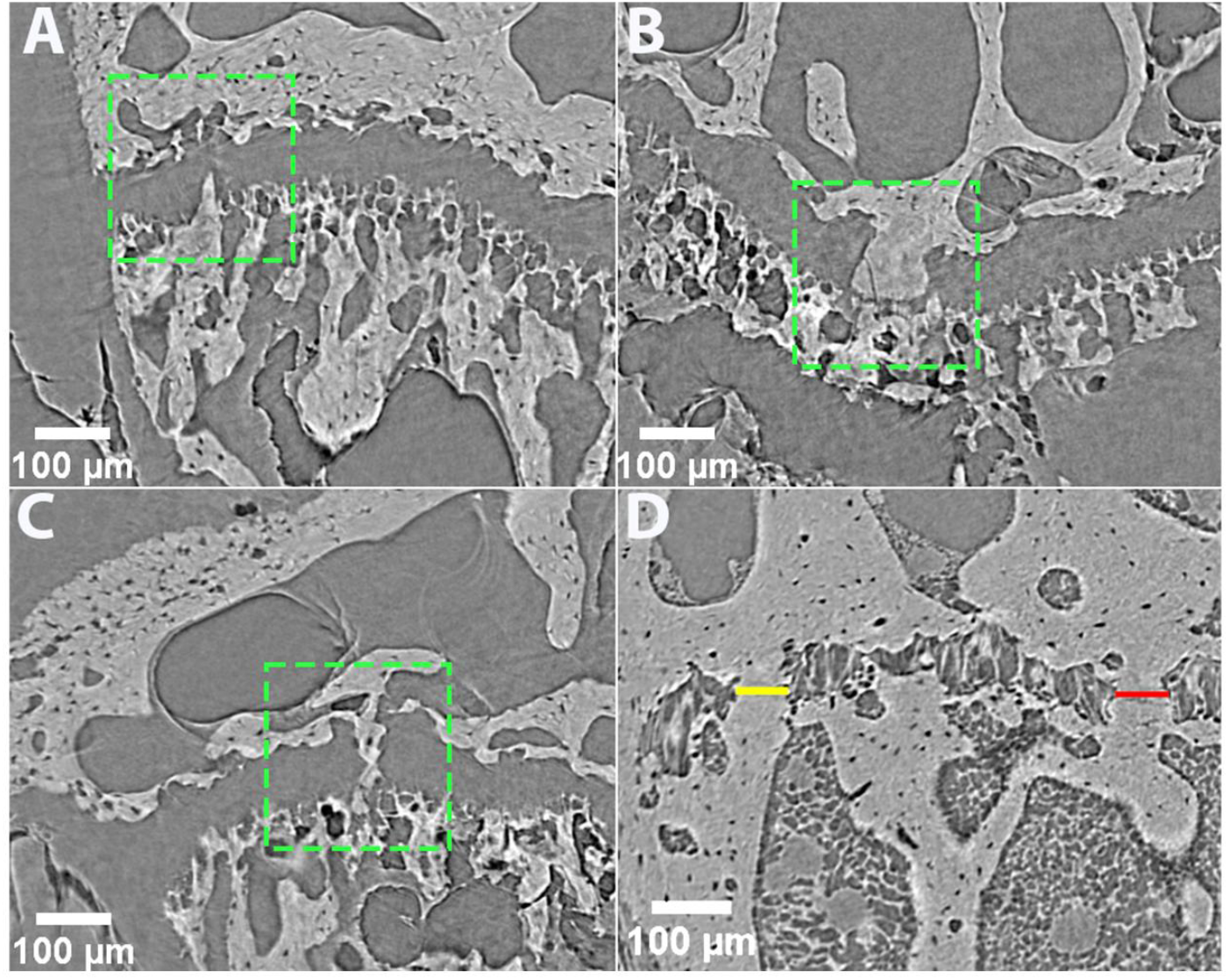
High-resolution sCT resolves GP bridges in detail. Dashed green squares highlight bridges that appear to be forming in a 10-week CBA mouse from the (A) metaphyseal and (B) epiphyseal GP surfaces, and (C) a mature bridge spanning the GP. Method of GP bridge measurement is shown in a 40-week CBA mouse; the bridge indicated with yellow has a width measured as 52.4 μm and the bridge indicated in red has a width of 68.3 μm.

It is evident that ex vivo imaging provides only meagre characterisation of bridge dynamics. To better explore the mechanisms that control bridge development and their temporal evolution, we chose to more fully probe the dynamics of their regulation by load using in vivo µCT scans obtained at several time-points in the same mice from a longitudinal 14-week-long study, incorporating two initial weeks of right limb loading. This confirmed a reduction in GP bridge density after a period of in vivo joint loading and extended this to show that the location of this reduction was almost entirely within the posterior, most loaded GP aspect (Fig. 5A-B) (19). Importantly, evaluation of scans taken 12 weeks after cessation of the 2 week-long loading period indicated a new accrual of reformed bridges in this posterior compartment (Fig. 5C). These data imply that bridges are transient, rapidly restorable structures that are regionally controlled by dynamic load stimuli.

**Figure 5.**
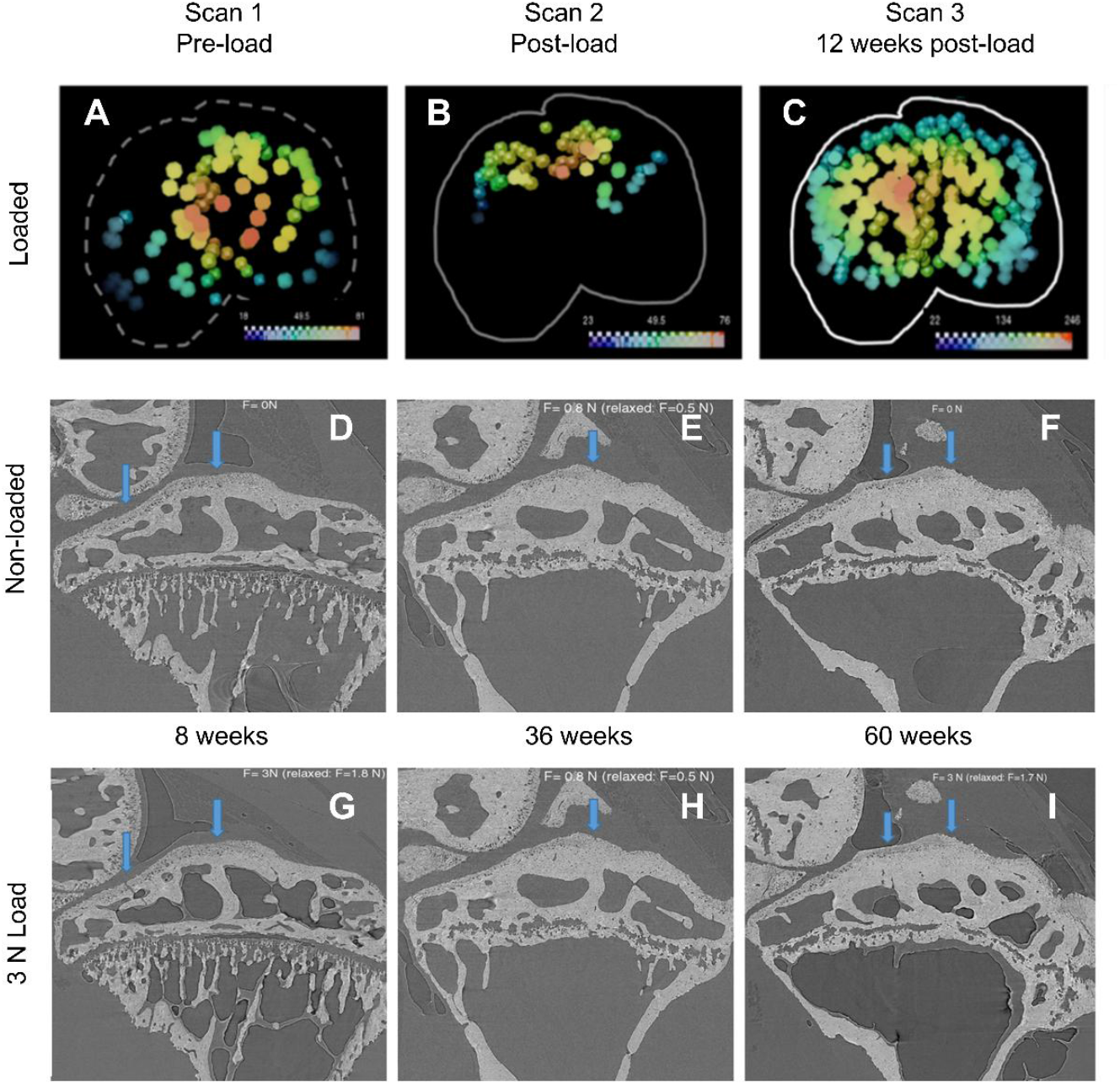
Bridge distribution and areal density at different time-points. Bridge number in 12 week-old female C57/Bl6 mice decreases after 2 weeks of load application (A-B) and increases 12 weeks thereafter load cessation (C). White outlines correspond to GP margins and colours reflect areal density. High resolution sCT scans of native (D-F) and in situ loaded (G-I) STR/Ort joints at 8-, 36- and 60-weeks of age.

### 3.4 Real-time synchrotron imaging during joint loading reveals a shift from GP deformation in growing mice to metaphyseal load-transfer via the epiphysis in maturity

The spanning of epiphyseal-to-metaphyseal bone regions makes it tempting to propose that bridges serve a mechanical role, but there is no current evidence for this. To explore this load transfer role, we acquired high resolution sCT scans of intact STR/Ort mouse joints during in situ loading at various ages (8, 36 and 60 weeks). This imaging confirmed an increase in bridge number with age, with ∼35 per GP at 8 weeks and ∼100 at 36 weeks of age (Fig 5D-F) and showed that load application resulted in discernible deformation of the cartilaginous GP in 8-week-old mice, discernible as a convergence of the bony epiphyseal and metaphyseal regions (Fig. 5D vs. G) but much less so in 36- or 60-week-old older mice (Fig. 5E vs. H and Fig. 5F vs. I). These low GP deformation levels in joints of older mice aligned with relative bridge abundance and load-induced metaphyseal bone displacement, showing that load-induced deformation in the young murine GP switches to metaphysis in mature mice [https://drive.google.com/file/d/17ei305pzCxj6mbFzPKVPB8F0QUPbbFEj/view?usp=sharing)].

### 3.5 DVC maps joint loading-induced inhomogeneities in GP strains which are coordinated to bridges

To probe local mechanics, load-induced in situ strains within the GP were estimated using DVC data from intact joints imaged by sCT in which <4 N loads were applied; this would also enable mapping to bridge location. Displacement point cloud surfaces were generated and tracked by DVC on both epiphyseal and metaphyseal sides of the GP. 3D linear quadratic brick finite elements were then defined to fill the space in between the surfaces. DVC measurements were mapped to element nodes, and strain values throughout the element interiors were calculated from shape function derivatives, generating cumulative strain tensor values within the GP. Strain plots showed that load-induced compression and tension were non-homogeneously distributed in the GP of an 8-week-old STR/Ort mouse (Fig. 6A-D), with maximum tensile (1^st^ principal) strain levels at a peak in the lateral/posterior side of the GP (Fig. 6B). DVC also revealed that joint loading generated greatest maximum shear on the lateral side of the GP, with a slight posterior predominance (Fig. 6C). In contrast, maximum compression (3^rd^ principal) was more homogeneously distributed with subtle anterior predominance, Fig. 6D). Overall, anterior/medial regions contained least GP strains and greatest load-induced strains were engendered in the posterior GP (Fig. 6A-D).

**Figure 6.**
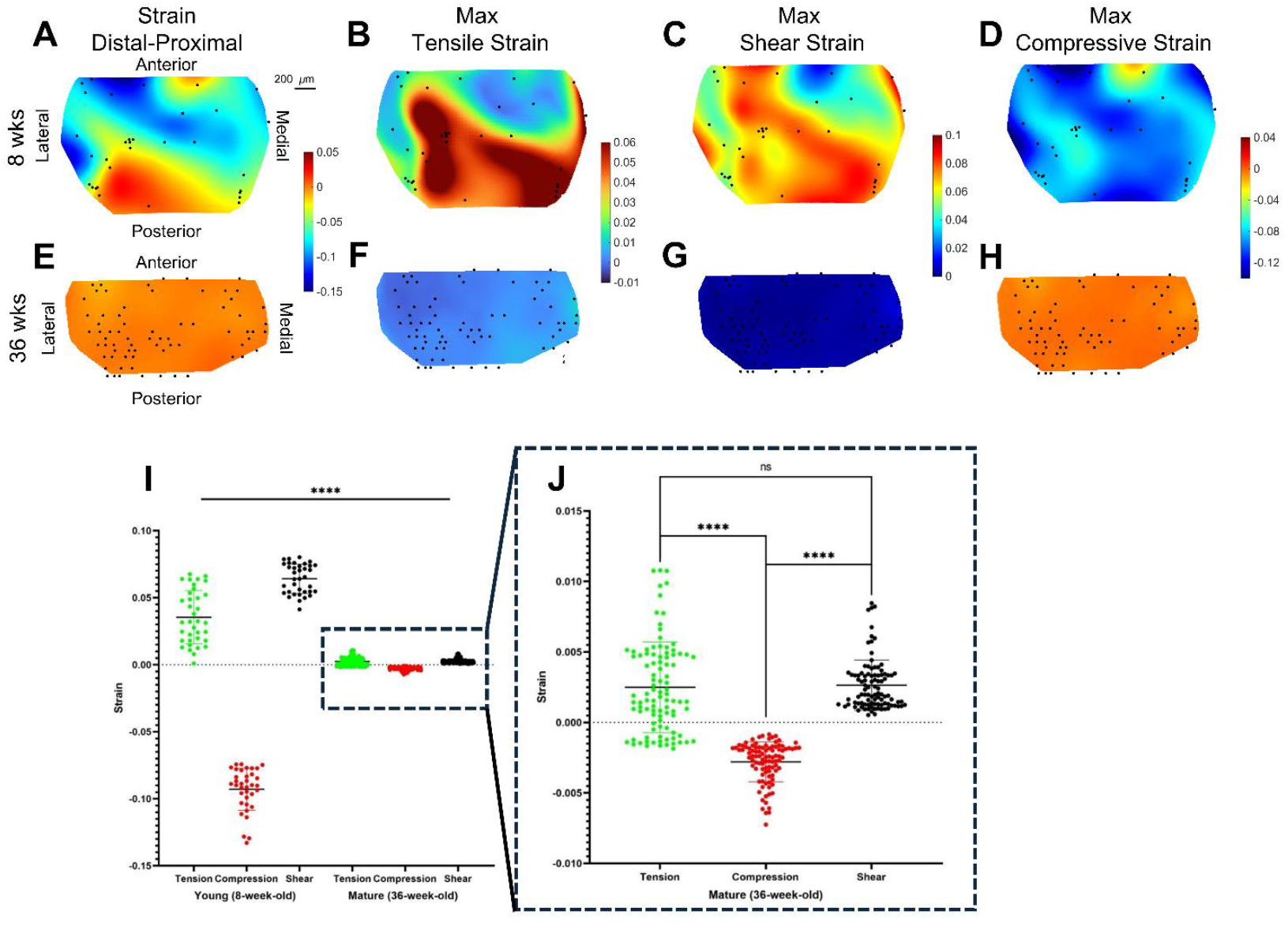
Colormaps of strain types engendered in GP of 8 week STR/Ort with static compressive load applied (A, total strain (top-bottom); B, tension, C; maximum shear and D; compression were measured and overlaid with bridge location (black dots) compared with similar in loaded 36 week STR/Ort mouse (E-G, respectively). Values at each bridge for tension, compression and shear strain in 8- and 36-week-old mice (I). At 8 weeks, significant difference is reached for tensional, compressive and maximum shear strain but in 36-week-old mice, significant difference is reached for tension vs. compression, shear vs. compression, but not for tension vs. shear. ****p<0.0001.

Local strains were also extracted from images of the 36-week-old joint in which skeletal growth will have halted (Fig. 6E-H). Use of DVC showed that both load-related compression and tension were markedly lower, with a narrower range (0.02 strain) than in the juvenile GP (Fig. 6E-H), indicating that GP deformation levels are lower in older mice. In contrast to the juvenile, strains in the 36-week-old GP were more homogeneous, with tensile stress showing the most marked range and highest values in medial-posterior, rather than in lateral regions. Mapping in the 36-week-old mouse showed that compressive (3^rd^ principal) strains in the GP were also more homogeneous (Fig. 6H). Loading of the 36-week-old joint induced greatest shear in the medial GP, lacking anterior/posterior dominance (Fig. 6G) indicating that greater, more inhomogeneous strains are engendered in young, growing mice.

To better determine the mechanical role of bridges, their location was superimposed onto the DVC colourmaps and corresponding strain values measured (Fig. 6I/J). This confirmed that magnitudes of load-induced compression, tension and shear strains generated at individual bridge locations was significantly greater in 8 than 36-week-old mice (p<0.0001, Fig. 6I/J); these parameters had similar relationships at both ages with compression that was around three times greater in magnitude than tension in the young GP, and almost twice the absolute magnitude of bridge-associated shear stress.

Interrogation of the 36-week-old mouse revealed that magnitude of bridge-associated compression was significantly greater than either tension or shear (p<0.0001); tensile and shear bridge-associated magnitudes were similar, with tension showing a somewhat wider range (Fig. 6J). These data also allow average bridge-associated strain magnitudes to be calculated in juvenile and mature mice. This revealed diminishing 14-, 21- and 31-fold lesser magnitudes of load-induced bridge tension, shear and compression, respectively in mature versus young GP (Supp. Table 3), supporting a role for bridges in primarily *absorbing* compression and shear applied across the GP.

### 3.6 Finite element modelling reveals that bridges serve to concentrate load-induced strain

To explore these proposed links between the presence of bridges in the GP and the local mechanisms of load transfer we also performed FE simulation. The image-based FE model of a representative proximal tibia specimen was developed to analyse the biomechanical effects of mineralised GP bridges (Fig. 7A) and the commonly used biomechanical parameter, von Mises stress distribution, in the epiphysis is shown in Fig. 7. The stress levels in the hyaline GP (Fig. 7B) differed by at least two orders of magnitude from those in the hard tissue, including in the opposing epiphyseal and metaphyseal bone plate and the mineralised bridges themselves (see Fig. 1C). A refined colour scale focusing on the hyaline GP (Fig. 7B) revealed that mineralised bridges locally increased stress levels in certain areas (black dashed circle), but not consistently across all regions (white dashed circle). On the other hand, the mineralised bridges consistently generated stress hotspots in the hard tissues (Fig. 7C), with stress levels reaching up to five orders of magnitude higher in these localised regions compared to non-bridged areas. These findings highlight the significant impact of bridges on the mechanical environment of the GP and surrounding tissues.

**Figure 7.**
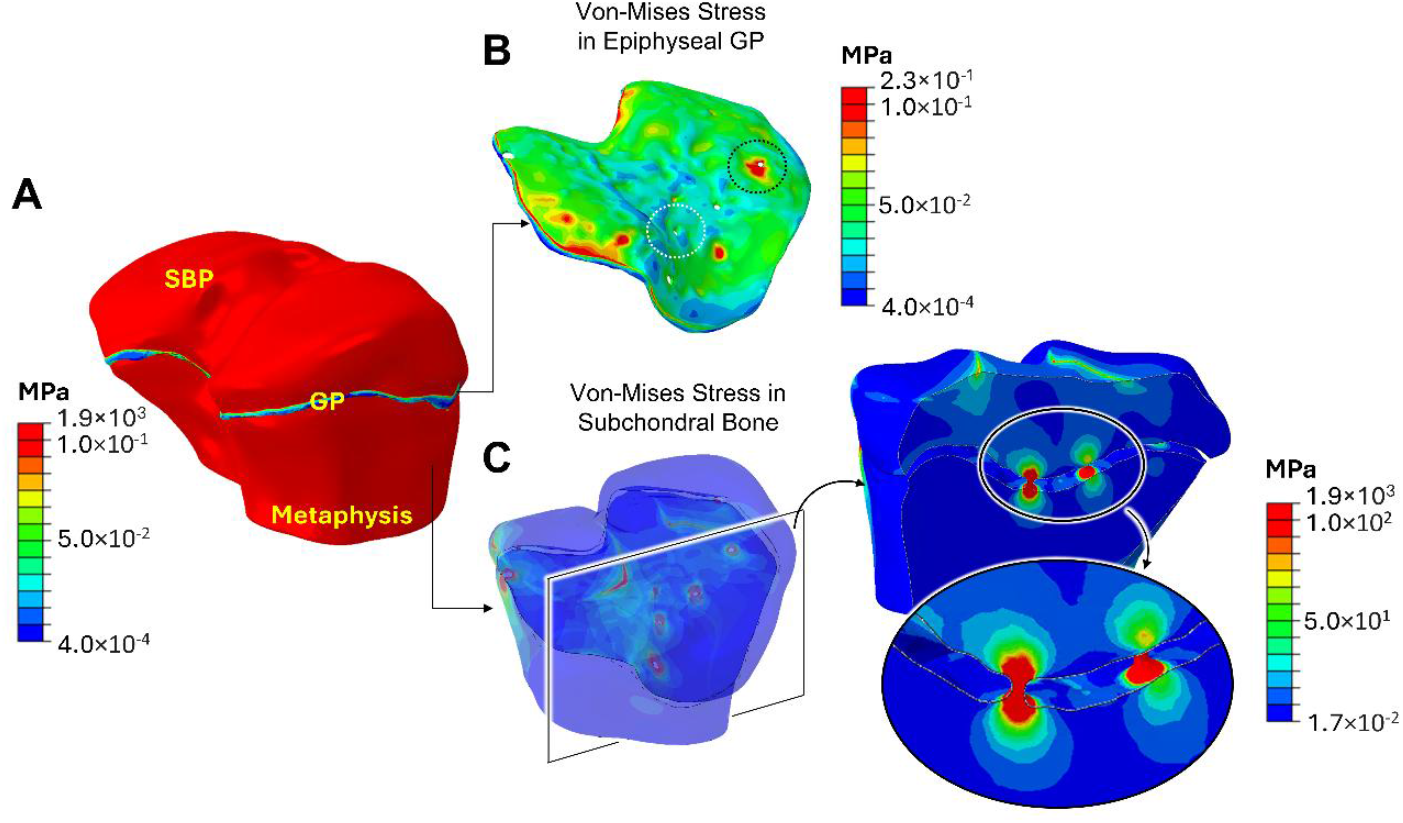
Biomechanical effects of bridge presence in the GP were assessed using von Mises stresses in the epiphysis through image-based finite element analysis (A). Von Mises stress distribution is shown for a representative specimen, ranging from 4.0×10^-4^ to 1.9×10^3^ MPa (B) with regions of bright red (high stresses) primarily appearing in the hard tissues. The GP is isolated and visualised with a refined stress range of 4.0×10^-4^ to 2.3×10^-1^ MPa, revealing the heterogeneity of stress distribution. Localised stress concentrations were observed around some bridges (e.g., marked by the black circle) but were not consistently present (e.g., as seen in the white circle). Mineralised tissue is isolated and visualised with a refined stress range (1.7×10^-2^ to 1.9×10^3^ MPa) highlighting stress hotspots at bridges; hotspots exhibited stress levels several orders of magnitude higher than in non-bridged areas (C).

### 3.7 Synchrotron imaging reveals consistent discontinuities in the mineral phase of GP bridges

Our data thus far suggest that bridge location is matched to distinct mechanical GP environments, but they do not explain how a mineralised epiphyseal-metaphyseal ‘bridge’ continuum facilitates further longitudinal GP expansion without generating a restraint upon tissue expansion. To interrogate this paradox further we performed more detailed examination of the mineral phase by histology and in µCT and sCT images. This, intriguingly, revealed that the bridges exhibited histological staining patterns that were more akin to surrounding non-calcified GP cartilage than local bone (Fig. 8) and CT images indicated the presence of a discontinuity in the mineral phase of all bridges, consistently found close their metaphyseal aspect (Fig. 4); these ‘gaps’ thus create an interruption to a presumed mineralised epiphyseal-metaphysis bridge continuum. These GP discontinuities were observed in both µCT or sCT scans and in GPs of immature and mature mice of different genetic backgrounds, thus challenging our view that these structures should be referred to as *tethers* or bone *bridges*.

**Figure 8.**
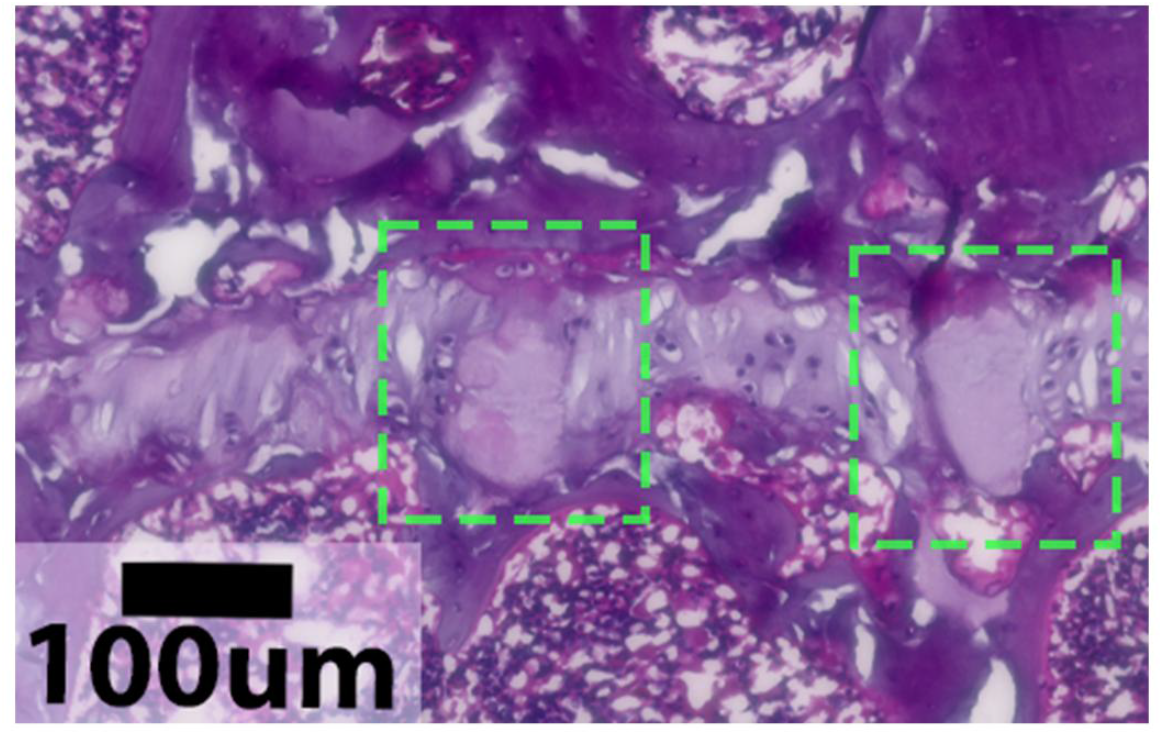
Bridges (dashed green) observed in H&E section of medial tibia in 36 week-old male CBA after sCT. Basophilic staining of bridges is more akin to surrounding non-calcified GP cartilage (pale lilac) than local bone (darker purple).

## 4. Discussion

Herein, we evaluated mouse GP bridge evolution with age, sex, pharmacological and biomechanical regulation, and have mapped bridge location to epiphyseal load-transfer using sCT and DVC in intact joints. Bridges have long been recognised, yet their function has remained elusive. We provide ground-breaking shifts in our knowledge of bridge function by revealing that they are dynamic, regulated, mechano-adapted structures, architecturally suited to a ‘base isolator’ role, absorbing compressive and shear forces that are transferred via the secondary ossification centre. Vitally, our data provide a new view on how bridges may allow dual GP roles, by enabling longitudinal growth and concurrently preserving mechanical stability of the epiphyses during articulation.

Our sCT imaging suggests that osteoblast-mediated ossification, with allied osteocyte entombment, is not responsible for GP bridge formation. Of course, we cannot rule out that bridges are constructed from an acellular or hypermineralised *dead* form of bone (45) but it is more likely that they are composed of calcified cartilage. Our data add the possibility that bridges are an early feature of all GPs, even those that will ultimately fuse fully to create a true physeal scar. Further, our observations suggest that bridges can originate from either GP face and can emerge or *disintegrate* by mechanisms that are seemingly unrelated to growth direction, suggesting they are transient structures which can be rapidly remodelled.

Consistent with earlier data, we find more bridges in ageing mice (9, 14, 29). It is an intriguing possibility that GP bridge number thus offer a novel and reliable method for defining mouse skeletal maturity; a stage that presently remains mostly speculative as it is often based on unsupported assumptions. Our data also reveal that the number of bridges depends not only on age but varies across mouse genotype as well. Given that bridges are found not just in mice but also pigs, rats and humans, it is evident that they are not randomly generated structures but likely linked to bone elongation.

Intermittent PTH delivered 5 days each week is known to generate cortical and trabecular bone accrual in mice and rats; our analyses in these same mice indeed confirm these PTH effects (20). It is surprising, therefore, that although this PTH regime also promotes longitudinal bone growth in rats (46), we find it does not modify bridges in mice. Earlier studies in rats found that GP thickness and longitudinal growth rate were both reduced by hindlimb unloading, and that both were reversed by PTH treatment in young but not adult rats (47). Our exploration of compartment-specific modifications by such intermittent PTH dosing in mice found bone accrual in all cortical and metaphyseal/epiphyseal trabecular compartments (20). Contrasting this to our observed lack of PTH effects on bridge number, further suggests that bridge formation is not mediated by osteoblast-mediated bone formation, but instead involves calcification of GP cartilage that is unaffected by PTH. This does not entirely align with data from PTHrP/PTH double-deficient mice, where PTH increases bone length and GP matrix mineralization (46). Thus, future studies may address how bridge number is affected by PTH in these and other mutant mice.

We also report sexual dimorphism in murine bridge number, with males possessing more than age-matched females. Based upon the assertion that large bridge number in mice is equivalent to GP closure in humans, these data may align to well-known pivotal roles for estrogen coincident with human sexual maturation. Conversely, whilst estrogen activity is needed for full pubertal development, GP maturation and closure, lower estrogen concentrations accelerate human growth and account at least partly for the pubertal growth peak both sexes (5, 20, 48). Indeed, estrogen-deficient individuals or those with estrogen receptor alpha (ERα) mutations exhibit a failure in GP fusion and slow but persistent growth into adulthood (49); similar growth persistence is also observed in ERα-deficient mice (50). Our data herein showing fewer bridges in females, implies that higher estrogen levels during growth may serve to restrict rather than promote bridge formation. Sexually dimorphic bridge regulation with hormonal links is endorsed by our data showing a reduction in bridges in female mice treated with tamoxifen; an anti-estrogen that functions as a partial estrogen-agonist in bone, which can cause systemic estrogen depletion. This aligns with studies where tamoxifen reduces longitudinal bone growth via chondrocyte apoptosis and systemic IGF-1 suppression (51). Our data support faster accumulation of bridges at late growth phases as elongation decelerates and their systemic hormonal regulation.

Recent preclinical studies have explored the impact of SFX-01 (stabilised natural sulforaphane) on bone, to find that it modifies DNA methylation patterns resulting in marked anti-osteoclastogenic effects (21, 22, 39, 43). Findings by Javaheri and co-workers support SFX-01’s osteotrophic roles that improve bone mass (22). Our data showing significantly greater bridge numbers in young, but not mature mice treated with SFX-01, link this pharmacological anti-resorptive to bridge accumulation, suggesting that normal bridge removal mechanisms rely on osteoclastic actions or that SFX-01’s antioxidant effects target other bridge remodelling processes.

Magnetic resonance imaging of human GPs during premature closure and trauma has disclosed large pathologically-linked bone bridges or *bars* (13, 52, 53). Evidently, these disrupt endochondral ossification and can consequently lead to skeletal deformity and growth arrest in children (54). Whether the GP bridges found in healthy animals are linked to these larger *bars* arising in response to mechanical injury is unknown, but deciphering the bridge formation mechanism will aid tissue engineering or regenerative strategies for treating dysfunctional GPs and in manipulating growth before adult height is attained.

There are many elegant studies and reviews showing GP behaviour can be mechanically modulated (55-59). This concept was indeed proposed many years ago in what is now known as the Hueter-Volkmann *principle* (60); that static compressive and tensional forces serve to restrict and promote GP-derived growth, respectively (55, 61). It is vital, however, to recognise that the GP habitually experiences dynamic gait-related mechanical loads during growth and that stability of neighbouring epiphyses is paramount. Our data show that despite bridge number being unaffected immediately post-loading in young-growing mice, that it is significantly reduced after six dynamic loading episodes in males and females over a two-week long period. It is an intriguing possibility that this load-induced reduction in bridge number may ultimately be connected to modified longitudinal bone growth. Earlier data have shown that the application of intermittent load can accelerate tibial lengthening (55).

It is also noteworthy that our data reveal rapid and reversible load-induced reduction in number and areal density of bridges in posterior epiphyseal regions. This aligns spatiotemporally with our earlier data using this model that reported that most applied loads are experienced in this region and endorses the view that bridging is not a permanent irreversible event (19). A further hypothesis is that bridges serve to tether longitudinal growth, restricting metaphyseal growth to a pre-determined speed. In line with this hypothesis, the reduced bridge number in loaded mice could therefore permit growth at an enhanced rate.

Direct strain measuring methods using gauges have previously been unable to quantify GP strains (58, 62). Our development of DVC analysis methods successfully overcomes this issue and have been applied herein to resolve strains that can be spatially-mapped to bridge locations. This is indeed the first report of 3D strain tracking and quantification in the GP within an intact limb under controlled and physiologically realistic loading conditions. Our DVC analysis shows how loading of intact epiphyses is translated into GP strain accumulation in young, growing mice and that the magnitude of these strains is markedly reduced upon attainment of skeletal maturity. Our mapping of engendered strain fields to existing bridge locations helps explain endochondral mechanics by showing that bridges serve as foci for highest compressive and shear strains in the young, which are diminished in mature mice. This supports a role for bridges in absorbing compression/shear to ensure mechanical GP functional competency. It is tempting to infer that load-induced mechanical targeting of the GP via these load-regulated bridges allows for faster/slower growth by removing/preserving bridges. Our FE analyses also align with this possibility. There is, however, need for a more refined FE modelling to include joint soft tissues – non-visualisable in our scans – to enable us to address how anatomical variations in GP bridges may affect mechanics of the whole epiphysis. All in all, these studies offer a better understanding of the GP behaviour during endochondral ossification and growth cessation.

Bridge fine structure, resolved by sCT imaging, reveals that discontinuities in their mineral phase is a consistent feature. This implies that the bridging process is controlled to uniformly conserve a more compliant, presumably cartilaginous interface which we have often found to be located close to the metaphyseal GP aspect. This sandwiching of a more compliant zone between two much stiffer, mineralised layers is remarkably similar to the ‘base isolation systems’ that architecturally decouple a highly stabilized superstructure (here, the epiphysis) from its substructure (here, the metaphysis). In building design, ‘seismic’ base isolation strategies are utilised in earthquake-prone regions to mitigate the superstructure response to an earthquake, effectively achieving isolation at the expense of deformations due to the lower lateral stiffness of the isolation system (63, 64). This is consistent with our observations of a conversion of applied compressive loading across the GP, into very high levels of shear force in the proximity of the bridges. It is compelling, to hypothesise that the conservation of a more compliant, presumably cartilaginous interface, within the bridge fine structure may therefore serve to facilitate interstitial GP growth for longitudinal bone expansion, whilst simultaneously contributing to preserving mechanical stability of the epiphysis during articulation. It is tempting to suppose that our description of bridge fine structure has uncovered how the GP balances these dual and complex functions across this challenging bone-cartilage-bone interface. Factors determining the specific localisation of bridge formation remain, however, unknown and studies are needed both to examine neighbouring matrix organisation and to track new bridge formation in vivo.

In conclusion, our results herein suggest that GP bridges are sex-, age-, strain-, and load-dependent. Our findings also highlight them as mechano-sensitive structures which may serve much like a ‘base isolator’. We propose that the function of these discontinuous structures is most pronounced during phases of slower longitudinal growth, during which they effectively separate the epiphysis from the metaphysis to both reduce the amount of energy that is transferred to the articular regions, whilst simultaneously engendering continued scope for lengthwise GP-mediated bone expansion.

## CRedIT authorship contribution statement

Conceptualisation: DV, KS, AAP; Methodology: DV, LAEE, CMD, AS, BJ, JC, RTH, MH, SM, PL, AB, BB; Software: DV, LAEE, CMD, AS, JC, MH, AB, BB; Formal analysis: DV, LAEE, CMD, JC; Resources: PDL, KS, and AAP; Writing – original draft: DV, AAP, LAEE, AS, KS; Writing – review and editing: all authors; Visualisation: DV, LAEE, AS; Supervision: AAP; Project administration: PDL, KS and AAP; Funding acquisition: PDL, KS and AAP.

## Acknowledgements

Laboratory facilities were provided by the Royal Veterinary College, the University of Brighton and the Research Complex at Harwell. We thank C. Reinhardt and L. Sinclair (University of Manchester at Harwell) for access to the Deben CT500 mechanical testing rig and E. Burke O’Leary, C. Berruyer and V. Schoeppler for their assistance during beamtime at the European Synchrotron Radiation Facility (proposal LS-3124). This work was supported by the European Union’s Horizon 2020 research and innovation programme under Marie Sklodowska-Curie grant agreement No. 721432 (AAP was also a recipient during this period of support from EPSRC, Osteoarthritis Technology Network Plus: a multidisciplinary approach to the prevention and treatment of osteoarthritis EP/N027264/1), by the UKRI MRC (MR/V033506/1), the BLAST Network (BB/W01825X/1), Royal Academy of Engineering (CiET 1819/10), the Chan Zuckerberg Initiative (CZIF2021-006424 and CZIF2022-316777) and partly by the Engineering and Physical Sciences Research Council (EPSRC) through the New Investigator Award (EP/T008059/1).

