## Supplementary file for "Regulation and function of trans-physeal growth plate bridges: evidence for a mechanical ‘base isolation’ role to minimise epiphyseal shear stress"

### Supplementary Material

Supplementary Table 1: Mouse details and protocol in each experimental group.

| Group (Figure) | Sex (M/F) | Age (weeks) | Limb (L/R) | Mouse strain | In vivo Treatment (Reference) |
|---|---|---|---|---|---|
| **2A** | M | 14 | L/R | C57Bl/6 | None |
| **2B** | M/F | 14 | R | C57Bl/6 | None |
| **2C** | M/F | 18-22 | R | C57Bl/6 | None |
| **2D** | M | 8<br>18-22<br>33+ | R | C57/Bl/6<br>CBA<br>STR/Ort | None |
| **2E** | M | 5<br>8-12<br>14<br>18-22<br>33+ | R | C57/Bl/6 | None |
| **2F/G** | M | 8<br>18-22<br>33+ | R | CBA<br>STR/Ort | None |
| **3A** | F | 15 | R | C57/Bl6 | iPTH [20] |
| **3B/C** | M | 7/12 | R | C57/Bl6 | Tamoxifen/SFX-01 [22, 23] |
| **3D** | M | 33+ | R | STR/Ort | SFX-01 [22] |
| **3E** | F | 15 | R | C57/Bl6 | iPTH [20]<br>iPTH+Load [18, 20] |
| **3F/G** | F | 14 | R | C57/Bl6 | Load [19] |
| **3H** | M | 14 | R | C57/Bl6 | Load [19] |

Supplementary Table 2: Bridge widths in the GP of 10-week-old CBA mouse.

| Bridge Number | Size (μm) |
|---|---|
| 1 | 28.3 |
| 2 | 14.8 |
| 3 | 14.4 |
| 4 | 9.9 |
| 5 | 43.2 |
| 6 | 42.8 |
| 7 | 27.1 |
| 8 | 14.7 |
| 9 | 8.0 |
| 10 | 20.9 |

Supplementary Table 3: Mean bridge strain values for tension, compression and shear in 8- and 36-week-old mice.

| | Tension | Compression | Maximum Shear |
|---|---|---|---|
| 8 week-old | 0.0355 | -0.093 | 0.064 |
| 36 week-old | 0.0025 | -0.003 | 0.003 |
| Fold change (young/mature) | 14x | 31x | 21x |
